# Single-cell multiomic analysis reveals the involvement of Type I interferon-responsive CD8+ T cells in amyloid beta-associated memory loss

**DOI:** 10.1101/2023.03.18.533293

**Authors:** Nilisha Fernando, Jaanam Gopalakrishnan, Adam Behensky, Lauren Reich, Chunhong Liu, Victor Bass, Michaella Bono, William Montgomery, Raffaella De Pace, Mary Mattapallil, Vijayaraj Nagarajan, Stephen Brooks, Dragan Maric, Rachel R Caspi, Dorian B. McGavern, Han-Yu Shih

## Abstract

Alzheimer’s disease (AD) is the leading cause of dementia worldwide, but there are limited therapeutic options and no current cure. While the involvement of microglia in AD has been highly appreciated, the role of other innate and adaptive immune cells remains largely unknown, partly due to their scarcity and heterogeneity. This study aimed to study non-microglial immune cells in wild type and AD-transgenic mouse brains across different ages. Our results uncovered the presence of a unique CD8+ T cell population that were selectively increased in aging AD mouse brains, here referred to as “disease-associated T cells (DATs)”. These DATs were found to express an elevated tissue-resident memory and Type I interferon-responsive gene signature. Further analysis of aged AD mouse brains showed that these CD8+ T cells were not present in peripheral or meningeal tissues. Preventing CD8+ T cell development in AD-transgenic mice via genetic deletion of beta-2 microglobulin (*B2m*) led to a reduction of amyloid-β plaque formation in aged mice, and improved memory in AD-transgenic mice as early as four months of age. The integration of transcriptomic and epigenomic profiles at the single-cell level revealed potential transcription factors that reshape the regulomes of CD8+ T cells. These findings highlight a critical role for DATs in the progression of AD and provide a new avenue for treatment.

## Introduction

Alzheimer’s disease (AD) is a debilitating neurodegenerative disease that leads to cognitive decline and memory impairment (Murphy and LeVine, 2010). The pathological hallmarks of AD include the formation of amyloid-beta (Aβ) plaques and neurofibrillary tangles in the brain (Hardy and Selkoe, 2002; Masters et al., 2015). Research in humans and mouse models have highlighted the crucial role of the immune system in neuroinflammation associated with these deposits (Chen and Holtzman, 2022; Robinson et al., 2017). While microglia, the most abundant macrophage-like immune cells in the central nervous system (CNS), have been studied extensively (Chen and Colonna, 2021; Leng and Edison, 2021; Salter and Stevens, 2017), the contribution of other immune cells, such as T cells and innate lymphocytes, in aging and neurodegeneration remains elusive (Chen and Colonna, 2022; Mason and McGavern, 2022).

T cells and innate lymphocytes are critical for modulating immune responses against pathogen invasion via the production of selective cytokines that have also been found to regulate neuronal activities. For instance, interferon-gamma (IFN-γ), a key cytokine involved in Type I immune responses during viral infection, plays an essential role in cognition and neuronal connectivity (Filiano et al., 2016). IFN-γ is specifically produced by lymphocytes including natural killer (NK) cells, Type I innate lymphoid cells (ILC1s), Type I helper CD4+ T (Th1) cells and cytotoxic CD8+ T cells. Type II cytokines, such as interleukins (IL)-4, IL-5 and IL-13, secreted by Th2 cells and ILC2s during helminth infection, play critical neuroprotective roles in homeostasis (Derecki et al., 2010; Quarta et al., 2020). Conversely, the Type III cytokine, IL-17, primarily expressed by Th17 cells, γδ T cells and ILC3s during fungal infection, can contribute to anxiety-like behaviors (Alves de Lima et al., 2020).

Emerging evidence suggests that peripheral lymphocytes may play critical roles in modulating brain aging and neurodegeneration via various mechanisms, such as cytokine production, plaque clearance, and intercellular communication (Castellani and Schwartz, 2020; Chen and Holtzman, 2022; Chen and Colonna, 2022; Mason and McGavern, 2022). In aging, NK cells accumulate within the neurogenic niche of the dentate gyrus, and their depletion through genetic manipulation or by antibody-based treatment has been shown to improve neurogenesis and cognitive function in both aging and AD models (Jin et al., 2021; Zhang et al., 2020). Conversely, ILC2s accumulate in the outer meninges and choroid plexus with age and have been associated with protection against cognitive decline (Fung et al., 2020). In *Rag2^-/-^Il2rg*^-/-^ AD mice, in which T, B and NK cell development are affected, reduced microglial phagocytosis was observed, which could be reversed with the addition of an immunoglobulin G (IgG) supplement, suggesting a role of IgG-producing B cells in neuroprotection (Marsh et al., 2016). Conversely, removing Foxp3+ regulatory CD4+ T cells mitigates AD-related pathology and reversed memory decline in 5xFAD mice, a transgenic (Tg) strain that expresses five human familial Alzheimer’s Disease (FAD) mutations (Baruch et al., 2015). A later study further showed that injection of Aβ-specific CD4+ T cells that secrete IFN-γ in the brains of 5xFAD mice enhanced Aβ clearance, while blocking CD4+ T cell development resulted in reduced microglial phagocytotic capacity and enhanced amyloid pathology (Mittal et al., 2019). On the other hand, presence of IL-17 producing γ8 T cells also contributes to cognitive impairment in a 3xTg-AD models (Brigas et al., 2021).

Tissue-resident memory CD8+ T cells have been observed to patrol the human brain parenchyma and perivascular spaces (Smolders et al., 2018). Recently, Gate et al. demonstrated that antigen-experienced, effector memory CD8+ T cells undergo clonal expansion within the cerebrospinal fluid (CSF) of human AD patients (Gate et al., 2020), while another group showed the infiltration of CD8+ T cells within the hippocampus in APP/PS1 transgenic mice to regulate neuronal-and synapse-related genes (Unger et al., 2020). Interestingly, depletion of CD8+ T cells in these mice with antibodies showed no effect on cognitive function or plaque deposition (Unger *et al*., 2020). Therefore, the role of CD8+ T cells in AD progression remains unclear.

Single cell and spatial transcriptomics approaches have been utilized to understand the contribution of disease-associated neurons and glial cells (including microglia, astrocytes, and oligodendrocytes) in the brains of human and mouse subjects (Kenigsbuch et al., 2022; Keren-Shaul et al., 2017; Mathys et al., 2019; Pandey et al., 2022; Silvin et al., 2022). A key finding was the identification of a TREM2-dependent disease-associated microglial (DAM) population that was increased in the 5xFAD mouse brain (Keren-Shaul *et al*., 2017). These DAMs were found to be in close proximity to Aβ plaques, with a higher expression of AD risk factors and genes involved in lysosomal and phagocytic pathways (Keren-Shaul *et al*., 2017). Supporting these findings, another spatial transcriptomics study using the *App^NL-G-F^* AD mouse model showed an upregulation of a plaque-induced gene network in microglia and astrocytes, involving complement, lysosomal and inflammatory pathways (Chen et al., 2020). Single nucleus RNA-seq profiling of human prefrontal cortex revealed the involvement of demyelination and inflammation in AD subjects (Mathys *et al*., 2019).

In the current study, we aim to uncover the role of lymphocytes in AD by dissecting lymphocyte transcriptomes and epigenomes in 5xFAD mouse brains, in which Aβ accumulation within neural tissues is a central pathological feature (Oakley et al., 2006). Our results revealed a disease-associated CD8+ T (DAT) cell population that is enriched in the brains of 5xFAD mice and exhibit tissue-resident, Type I IFN-responding signatures. Integration of transcriptomic and epigenomic profiles demonstrated that the alteration of AD-transgenic CD8+ T cell regulomes were shaped by transcription factors induced by IFN signals, such as interferon regulatory factors (IRFs). Imaging of brain sections from 5xFAD mice showed that CD8+ T cells deeply infiltrated the parenchyma and were found in proximity to neurons and microglial dendrites. These CD8+ T cells accumulated not only in hippocampus, but also in cortex and thalamus. Importantly, interference with CD8+ T cell development via genetic deletion of *B2m*, or removal of the IFN-alpha receptor (*Ifnar1*^-/-^) rescued memory decline in 5xFAD mice. Immunohistochemically, aged 5xFAD-*B2m*^-/-^ brains had significantly fewer Aβ plaques than the control 5xFAD brains. We propose that DATs play an essential role in AD pathogenesis through an IFN-dependent axis, and potentially offer new therapeutic opportunities.

## Results

### Lymphocytes are enriched in aging murine AD brains

Although extensive research has demonstrated the pivotal role of microglia in the clearance of Aβ plaques and dead cells during AD, the precise contribution of other immune cells remains an important area of investigation (Chen and Holtzman, 2022; Chen and Colonna, 2022; Mason and McGavern, 2022). We identified the role of non-microglial immune cells in a mouse model of AD by isolating CD45^hi^ cells from the brains of wild type (WT) and 5xFAD (AD) mice, a double transgenic model that carries five AD familial mutations for amyloid precursor protein (APP) and presenilins (PSEN1) (Oakley *et al*., 2006), at 1-2 months, 4.5-6 months and 10-14 months of age (Figure 1A). We performed single-cell RNA-sequencing (scRNA-seq, 10x Genomics) with cell hashing (n=4-6 per group). This sorting strategy allows profiling of all immune cells except microglia that express CD45 at an intermediate level. Unsupervised graph-based clustering by Seurat designated these non-microglial immune cells into six major groups: myeloid cells (including macrophages, neutrophils, dendritic cells, and microglial contamination), αβ T cells (including CD4+ and CD8+ T cells), NK cells/ILC1s, γδ T cells, B cells, and ILC2s, based on cell identity genes (Figure 1B-C and Supplementary Figure 1A). Comparing the proportions of the major cell groups revealed comparable composition of non-microglial immune cells in young (1-2 months) WT and AD mouse brains (Figure 1D and Supplementary Figure 1B). However, in AD mouse brains at 4.5-6 months of age, αβ T cells comprised close to 50% of all CD45^hi^ cells, a notable increase compared with less than 20% in WT brains. This was accompanied by a decrease in myeloid cells and γ8 T cells at 4.5-6 months of age. At 10-14 months, both WT and AD mouse brains showed a large increase in αβ T cells, alongside a decreased proportion of myeloid cells. This was even more prominent in the AD mice, where αβ T cells occupied over 60% of CD45^hi^ cells. Notably, a majority of the αβ T cells were derived from CD8+ T cells (Figure 1C and 1E).

**Figure 1.**
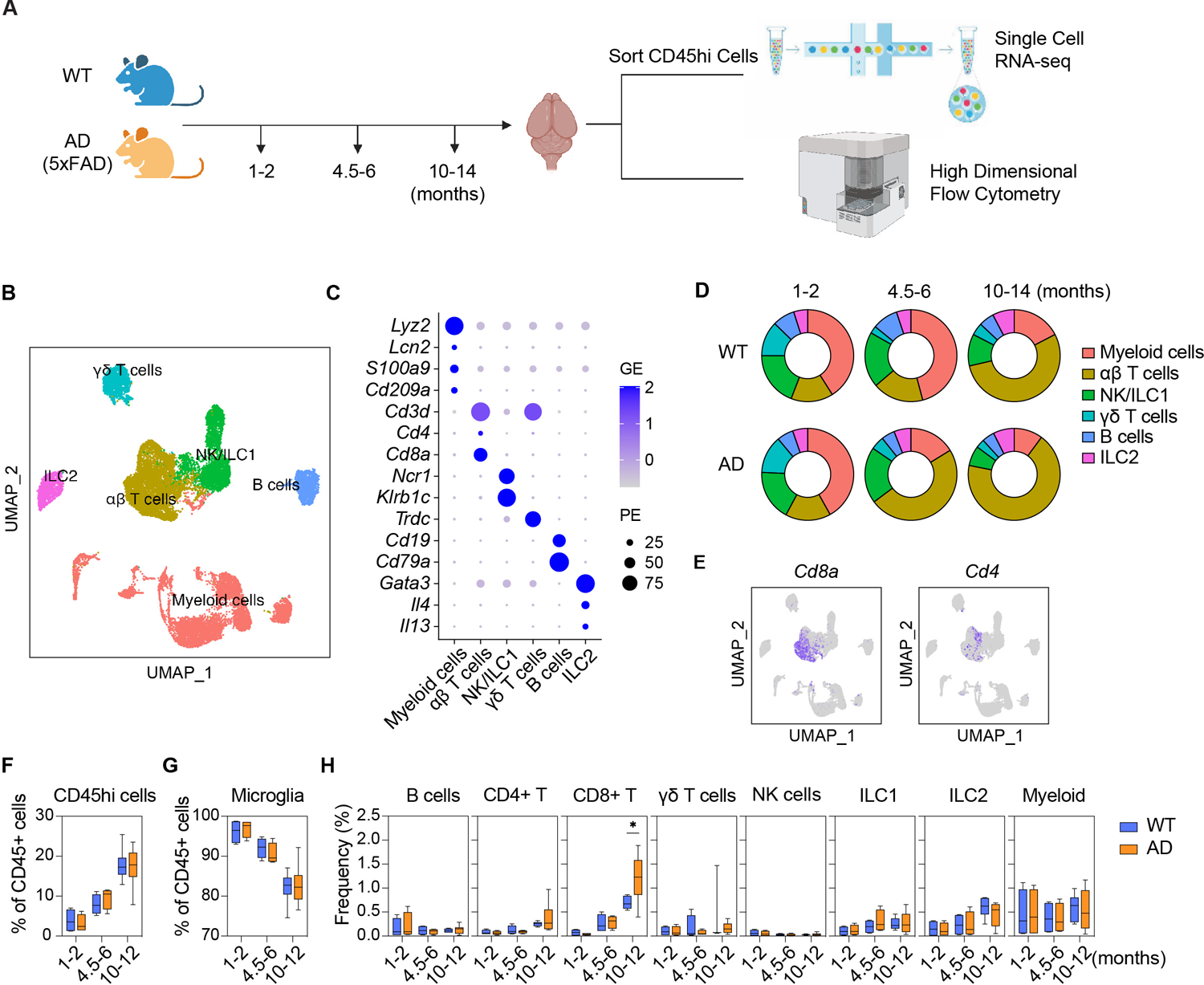
Enrichment of CD8+ T cells in aged AD mouse brains is revealed by profiling non-microglial immune cells. A. Experimental pipeline for analysis of wild type (WT) and 5xFAD (AD) brains at 1-2, 4.5-6 and 10-14 months of age. CD45^hi^ cells were either sorted for single cell analysis or were used for high-dimensional flow cytometry analysis. B. scRNA-seq UMAP of 20,874 cells combined between all experimental groups (n=4-6 pooled mice per group) showing myeloid cells, αβ T cells, NK/ILC1s, γδ T cells, B cells and ILC2s. 20,874 cells were pooled from WT 1-2 months (5,014 cells), AD 1-2 months (4,505 cells), WT 4.5-6 months (3,829 cells), AD 4.5-6 months (3,731 cells), WT 10-14 months (1,677 cells) and AD 10-14 months (2,118 cells). C. Dot plot showing lineage genes from the 6 cell clusters. D. Average cell proportions of the 6 major cell clusters between WT and AD mice at the 3 time points. E. Feature plots showing expression of selected genes (*Cd8a* and *Cd4*). F-H. Flow cytometry analysis (n=4-8) showing the frequency (%) per event of CD45^hi^ cells (B cells, CD4+ T cells, CD8+ T cells, γδ T cells, NK cells, ILC1s, ILC2s and myeloid cells) and microglia (CD45^int^ cells). A significant increase in CD8+ T cells in AD mice was found compared with WT at 10-12 months of age (p<0.05). Statistical analysis was performed using unpaired Student’s t tests. * denotes significance of p<0.05.

To validate the increase in the lymphoid compartment during neurodegeneration, we then performed high-dimensional flow cytometry (HDFC) on cells isolated from WT and AD mouse brains (n=4-8 per group) at three different ages for immune profiling (gating strategy shown in Supplementary Figure 1C). Although the proportion of CD45^hi^ cells increased and microglia (CD45^intermediate^) decreased with age, there was no significant difference between WT and AD mouse brains (Figure 1F-G). Profiling major populations including B cells, CD4+ T cells, CD8+ T cells, γδ T cells, NK cells, ILC1s, ILC2s and myeloid cells initially revealed no differences between WT and AD; however, at 10-12 months, the frequency of CD8+ T cells became significantly increased in AD mouse brains (Figure 1H).

Next, we investigated whether the enrichment of adaptive lymphocytes in AD mice was model-specific or due to APP overexpression. To exclude this possibility, we analyzed the hAbeta^SAA^ knock-in (APP-SAA KI) mice, which introduced humanizing Aβ mutations to endogenous exons without disturbing intrinsic gene expression (Xia et al., 2022). We profiled the CD45^hi^ cells in APP^SAA^ KI mouse brains at 8-9 months (n=3). Consistent with our prior observation, we observed a significant increase in CD8+ T cells in APP^SAA^ KI AD brains compared with WT (Supplementary Figure 1D). We also observed a significant increase in γδ T cells in APP^SAA^ KI AD brains, even though the frequency was much lower than CD8+ T cells. Taken together, the results from HDFC demonstrated a selective increase of CD8+ T cells in brains from two different strains of aged AD mice, which corroborates our scRNA-seq data.

## Brain infiltrating CD8+ T cells localize near neurons and microglia during AD

We further investigated the localization of lymphocytes within the AD mouse brains using multiplexing immunohistochemistry (IHC). In coronal sections containing the motor cortex and hippocampus regions from 10-14-month-old mice, we found a significant increase in CD8+ T cells in the parenchyma of AD mouse brains compared with WT brains (Figure 2A and Supplementary Figure 2A-B, n=2-3), including cortex, thalamus and hippocampus. In contrast, few CD4+ T cells were identified in WT and AD mouse brains, and few CD8+ T cells were identified in the WT brains, many of which were localized close to the leptomeninges (Figure 2A).

**Figure 2.**
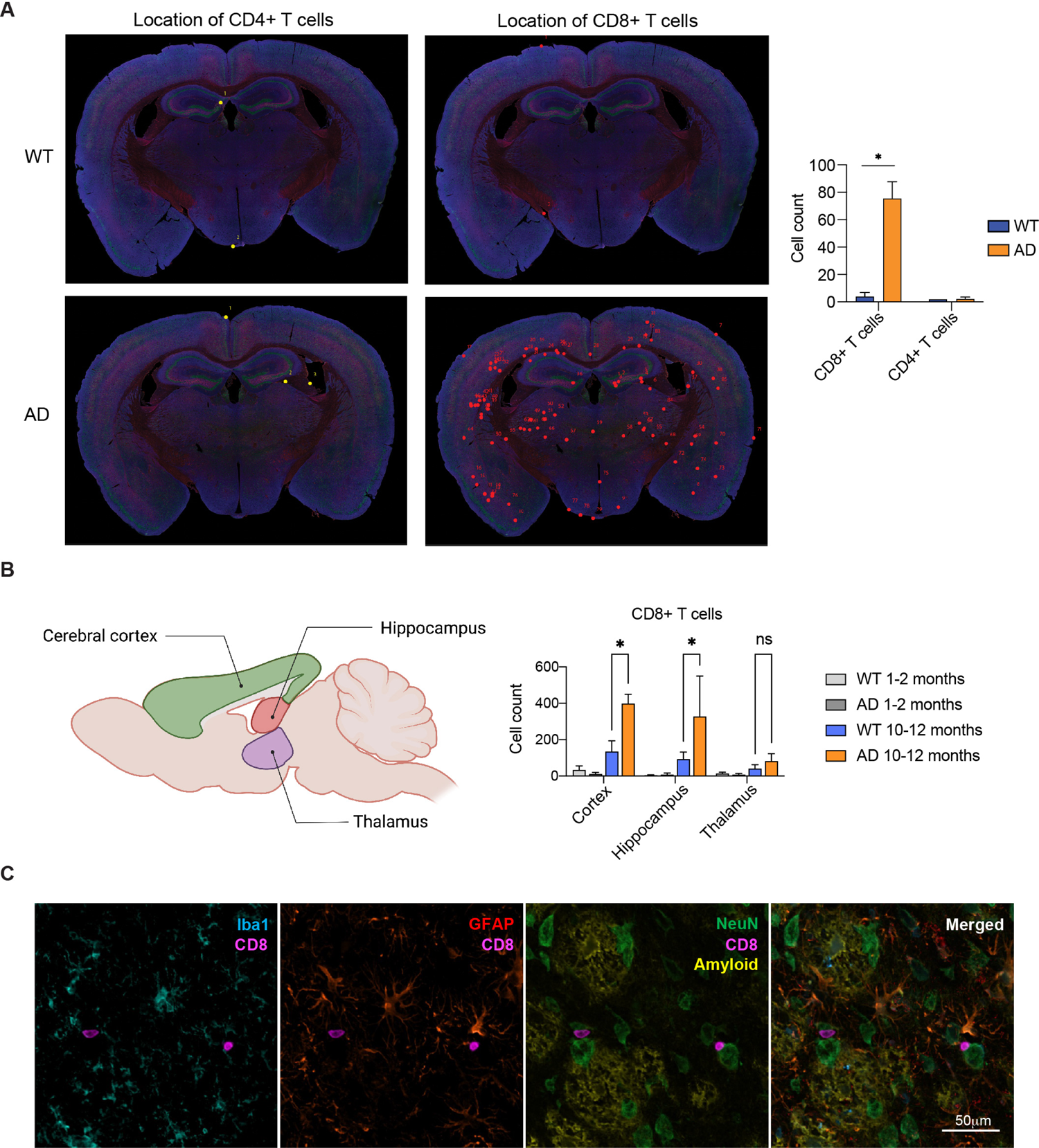
CD8+ T cells infiltrate the cortex and hippocampus of AD mouse brains. A. Multiplexing immunohistochemistry images depicting the location of CD4+ T cells (yellow) and CD8+ T cells (red) in wild type (WT, n=2) and 5xFAD (AD, n=3) brains. The quantitative measurements per brain slice is shown in the right panel. B. Flow cytometry analysis of CD8+ T cells in the cerebral cortex, hippocampus and thalamus of WT and AD mouse brains at 1-2 months and 10-11 months of age. C. CD8+ T cells (pink) in AD mouse brains shown near IBA1+ microglia/macrophages (blue), GFAP+ astrocytes (red), NeuN+ neurons (green) and amyloid (yellow). Scale bar is 50 µm. Statistical analysis was performed using unpaired Student’s t tests. * denotes significance of p<0.05.

We next performed HDFC to profile immune cells in the cerebral cortex, hippocampus and thalamus regions of the brain in young (1-2 months) and aged (10-11 months) mice. We found a significant increase in the number of CD45^hi^ cells in the aged AD mouse hippocampus (Supplementary Figure 2C). Further, CD8+ T cells were significantly more numerous in both the cerebral cortex and hippocampus in aged AD mouse brains, but not in young AD brains (Figure 2B). In contrast, no changes were observed in B cells or CD4+ T cells in either young or aged mice (Supplementary Figure 2C).

To gain a deeper understanding of the relationships of CD8+ T cells with other cells, as well as the pathological hallmarks in AD mouse brains, we also stained brain sections with antibodies against IBA1 (microglia/macrophages), NeuN (neurons), GFAP (astrocytes) and Aβ in our staining panel (Figure 2C). Our results revealed that CD8+ T cells in AD mouse brains were often juxtaposed to neurons or microglial processes, suggesting that CD8+ T cells might have the ability to affect these two brain resident cell populations.

## Single-cell analysis of lymphocytes reveals a unique ISG+ CD8+ T cell cluster in AD mouse brains

To enhance the clustering resolution, we next re-clustered the lymphocyte populations after excluding the myeloid cells (Figure 3A). Our analysis clearly differentiated individual clusters of NK cells, ILC1s, ILC2s, ILC3s, and γδ T cells, as well as two clusters of B cells. However, CD8+ T cells exhibited three sub-populations, including a general cluster, an interferon-stimulated gene (ISG+) expressed cluster, and a cytotoxic cluster based on their transcriptomic profile. The ISG+ CD8+ T cells expressed high levels of ISGs including *Ifi213*, *Iigp1*, *Ifit1*, *Ifit3*, *Igtp, Ifi206*, *Ifitm3*, *Isg20*, and *Slfn8* (Supplementary Figure 3A)*. Irf7*, a transcription factor regulating the expression of Type I IFN genes, was also expressed in ISG+ CD8+ T cells. Interestingly, *Ifng*, a Type II IFN, was expressed by all three CD8+ T cell clusters. The cytotoxic CD8+ T cell cluster, however, expressed higher levels of *Gzma*, *Gzmb*, *Zeb2* and *Klrg1*. Upon comparing cell proportions, we observed minor changes in young (1-2 month old) brains (Figure 3B). However, both general and ISG+ CD8+ T clusters increased in AD mouse brains at an intermediate age (4.5-6 months), with ISG+ CD8+ T cells remaining elevated at 10-14 months of age. The differences in cell number proportions were statistically significant as determined using a single cell proportion test (scProportionTest, Supplementary Figure 3B). Of note, while CD8+ T cells were significantly enriched in the AD mouse brains (Figure 3B and Supplementary Figure 3B-C), myeloid cells were not enriched after re-clustering (Supplementary Figure 3D-E).

**Figure 3.**
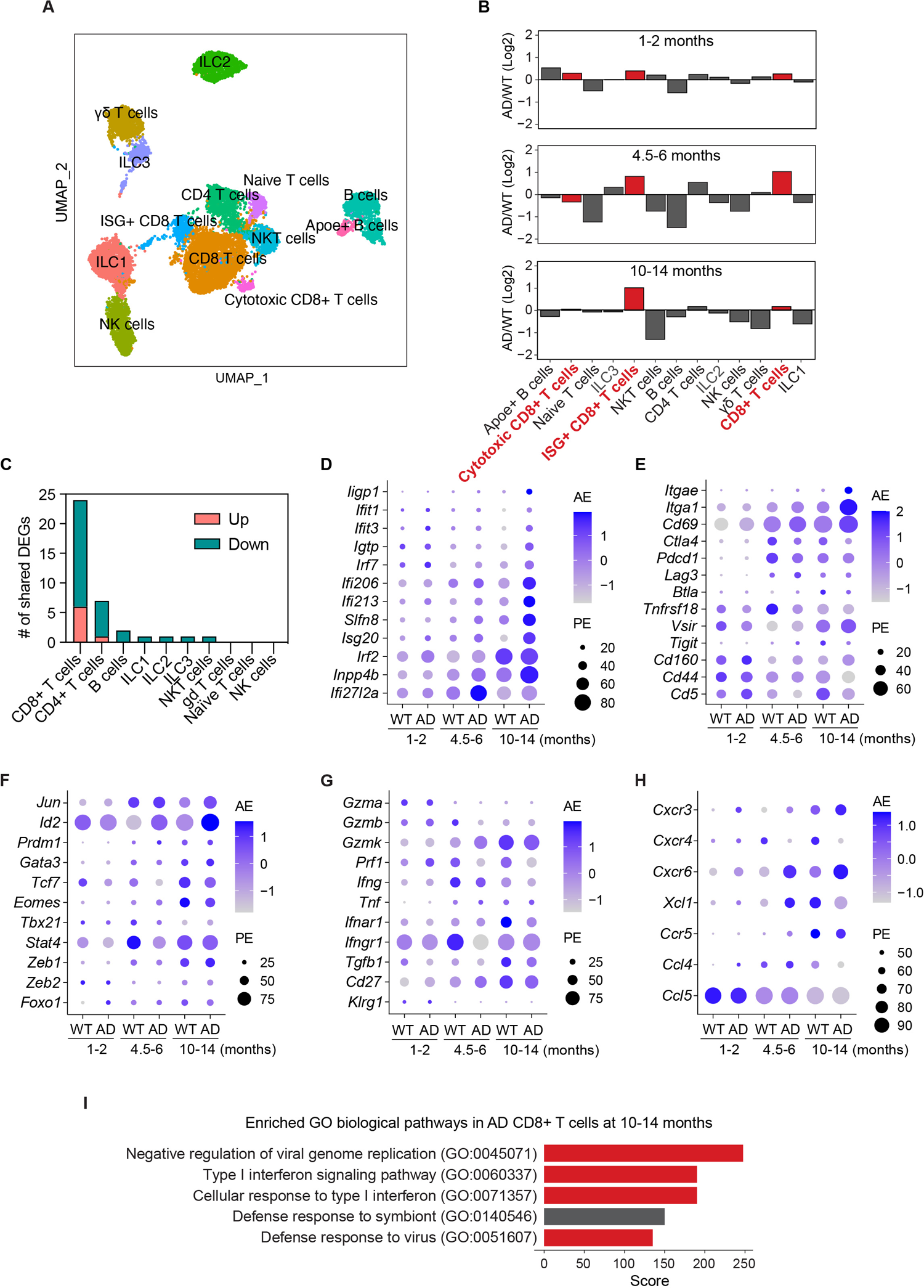
Single cell analysis of lymphocytes shows an increased Type I interferon (IFN) signature in CD8+ T cells from aging AD mouse brains. A. scRNA-seq UMAP of lymphocytes only depicting 13 cell clusters from wild type (WT) and 5xFAD (AD) brains at 1-2, 4.5-6 and 10-14 months of age (sample sizes provided in the Figure 1 legend). B. Cell proportion graphs per time point showing log2 fold change of AD compared with WT. CD8+ T cell clusters are shown in red. C. Number of shared differentially expressed genes (DEGs) between 4.5 months and 10-14 months per lineage in AD mice. D-H. Dot plots showing CD8+ T cell gene expression of Type I IFN-responding genes (D), residency and exhaustion markers (E), transcription factors (F), effectors (G) and chemokines (H). I. Enrichr gene ontology (GO) biological pathway analysis (cut off set at combined score >100 and adjusted p<0.05) on combined CD8+ T cells from AD mouse brains at 10-14 months showing the top 5 upregulated pathways.

Subsequently, we conducted differential gene expression analysis by comparing WT and AD mice for each cell lineage, including combined CD8+ T cell clusters and B cell clusters (Supplementary Figure 3F). A full listing of differentially expressed genes (DEGs) per lymphocyte lineage is included in Supplementary Table 1. As expected, there were fewer DEGs across all lineages at 1-2 months. At 4.5-6 months, several lymphocyte lineages, including NKT cells, ILC2s, CD8+ T cells and CD4+ T cells, revealed over 50 DEGs. However, at 10-14 months, CD8+ T cells appeared to be the predominant lineage that expressed dramatic transcriptomic changes. Specifically, 43 genes were upregulated, and 71 genes were downregulated in the CD8+ T cells from aged AD mouse brains, compared to age-matched WT. CD4+ T cells also showed many DEGs in aged AD mouse brains (Supplementary Figure 3F, Supplementary Table 1). We further compared DEGs at 10-14 months with DEGs at 4.5 months (Figure 3C and Supplementary Table 1) and found that CD8+ T cells were the lineage that shared the most DEGs (24 genes) between two time points, followed by CD4+ T cells (7 genes). The six upregulated DEGs shared between the two temporal CD8+ T cell populations are *Cxcr6*, *Ly6a*, *Inpp4b*, *Id2*, *Clec2d*, and a long-noncoding RNA (lncRNA) *BE692007*. The downregulated DEGs include, but are not limited to, transcription factors, *Zfp36l2, Junb, Fosl2, Crem,* and *Nr4a1*; dual specificity protein phosphatases, *Dusp1* and *Dusp5*; G-protein coupled receptors and relevant molecules, *P2ry10, Rgs2* and *Gpr183*; splicing factors, *Tra2b* and *Srsf5*; TNFα-induced protein *Tnfaip3*; and a lncRNA *Gm10076*. Interestingly, this lncRNA was also downregulated in other lineages such as CD4+ T cells, B cells, ILC1, ILC2, ILC3, and NKT cells. The role of this lncRNA remains to be investigated.

When combining all CD8+ T cell clusters for analysis, the majority of upregulated genes in aging AD mice were ISGs, including *Ifi206*, *Ifi213*, *Ifit1*, *Ifit3*, *Ifitm3*, *Isg20*, *Irf7*, *Iigp1*, *Igtp*, *Slfn8*, and *Inpp4b* (Figure 3D). We also found that some transcription factors (*Jun*, *Id2, Prdm1, Gata3, Tcf7, Eomes, Stat4*), effectors (*Gzmk, Prf1, Ifng, Tnf, Ifnar1, Tgfb1*), tissue-residency genes (*Itgae* (encoding CD103), *Itga1* (encoding CD49a) and *Cd69*), exhaustion/activation genes (*Ctla4*, *Pdcd1, Lag3, Btla*) and chemokines/receptors (*Cxcr3, Cxcr4*, *Cxcr6*, *Xcl1*, *Ccr5*) were upregulated with age (Figure 3E-H). Notably, canonical genes such as *Itgae, Itga1* and *Cd69* were significantly increased in AD CD8+ T cells, indicating that they represent tissue-resident CD8+ T cells (Figure 3E). Surprisingly, key CD8+ T cell effector genes such as *Prf1* and *Ifng* were downregulated in CD8+ T cells in aged AD mice. Conversely, *Gzma*, *Gzmb*, *Klrg1*, *Cd160*, and *Ccl5* decreased in expression with age, with minimal difference between WT and AD. Another interesting finding was that CD8+ T cells from AD mice exhibit upregulation of *Cxcr6*, a gene encoding the C-X-C motif chemokine receptor 6 (CXCR6), and downregulation of another chemokine receptor *Cxcr4* across all time points, when compared to WT. The chemokine receptor expression shifting from *Cxcr4* to *Cxcr6* recapitulates the T cell phenotype recently observed in human CSF associated with cognitive impairment (Piehl et al., 2022). Resident memory T cells that express CXCR6 are known to infiltrate tissues or tumors for immunosurveillance via interactions with its ligand CXCL16 expressed on certain myeloid cell types (Mabrouk et al., 2022). Our data support recent findings on the role of the CXCR6/CXCL16 axis in T cell recruitment in the brain during murine viral infection and human neurodegeneration (Piehl *et al*., 2022; Rosen et al., 2022) and also suggests that chemokine-mediated invasion of T cells and immunosurveillance may occur early in AD progression.

We next performed Enrichr gene ontology (GO) biological pathway analysis on the DEGs of CD8+ T cells. As expected, we found that anti-viral signaling pathways were upregulated, including Type I IFN signaling in AD CD8+ T cells (Figure 3I), consistent with recent findings using bulk RNA-seq approaches (Altendorfer et al., 2022). To our surprise, no other lymphocyte lineages revealed a clear anti-viral signature (DEGs in Supplementary Table 1). Recently, several studies have indicated Type I IFN signaling in glial cells in neurodegenerative brains, including astrocytes (Habib et al., 2020), oligodendrocytes (Kenigsbuch *et al*., 2022; Pandey *et al*., 2022) and microglia (Roy et al., 2022), suggesting that other non-lymphocyte cell types may also participate in Type I IFN signaling in the brain. Taken together, our data demonstrate a tissue-resident ISG+ CD8+ T cell cluster in aging AD mouse brains that may be first evident at 4.5-6 months of age, around the same time as early cognitive deficits are detected in this model (Schneider et al., 2014).

## Single-cell multiomic analysis of lymphocytes shows distinct regulomes in CD8+ T cells from AD mouse brains

To investigate how immune cells were regulated in AD, we next performed multiomic analysis (10x Genomics) to simultaneously measure the changes in transcripts (scRNA-seq) and chromatin accessibility (scATAC-seq) within a single cell in CD45^hi^ cells isolated from WT and AD mouse brains at 10-14 months of age (Figure 4A). Unsupervised graph-based clustering on nuclear RNA-seq data by Seurat revealed similar immune cell populations (Figure 4B-C) as previously identified using the regular scRNA-seq method (Figure 1B and Supplementary Figure 1A). However, when we combined scRNA-seq and scATAC-seq profiles for clustering, we found that WT and AD mice clearly differed in their clustering for CD8+ T cells, NK/ILC1, microglia, monocytes, and Ace+ macrophages between WT and AD mice; however, this was not the case for γδ T cells, ILC2s, B cells, dendritic cells, BAM, Mgl2+ and Clec9a+ macrophages (Figure 4D). We next asked whether the alteration of chromatin landscapes contributes to the high DEG number in CD8+ T cells. Among CD8+ T cell DEGs that were upregulated, *Id2* and the IFN-responsive genes, *Ifit1* and *Igtp*, displayed greater chromatin accessibility in AD mouse brains (Figure 4E). In contrast, *Cxcr4*, a gene that is downregulated in AD CD8+ T cells (Figure 3G), demonstrated a reduction of openness (Figure 4E).

**Figure 4.**
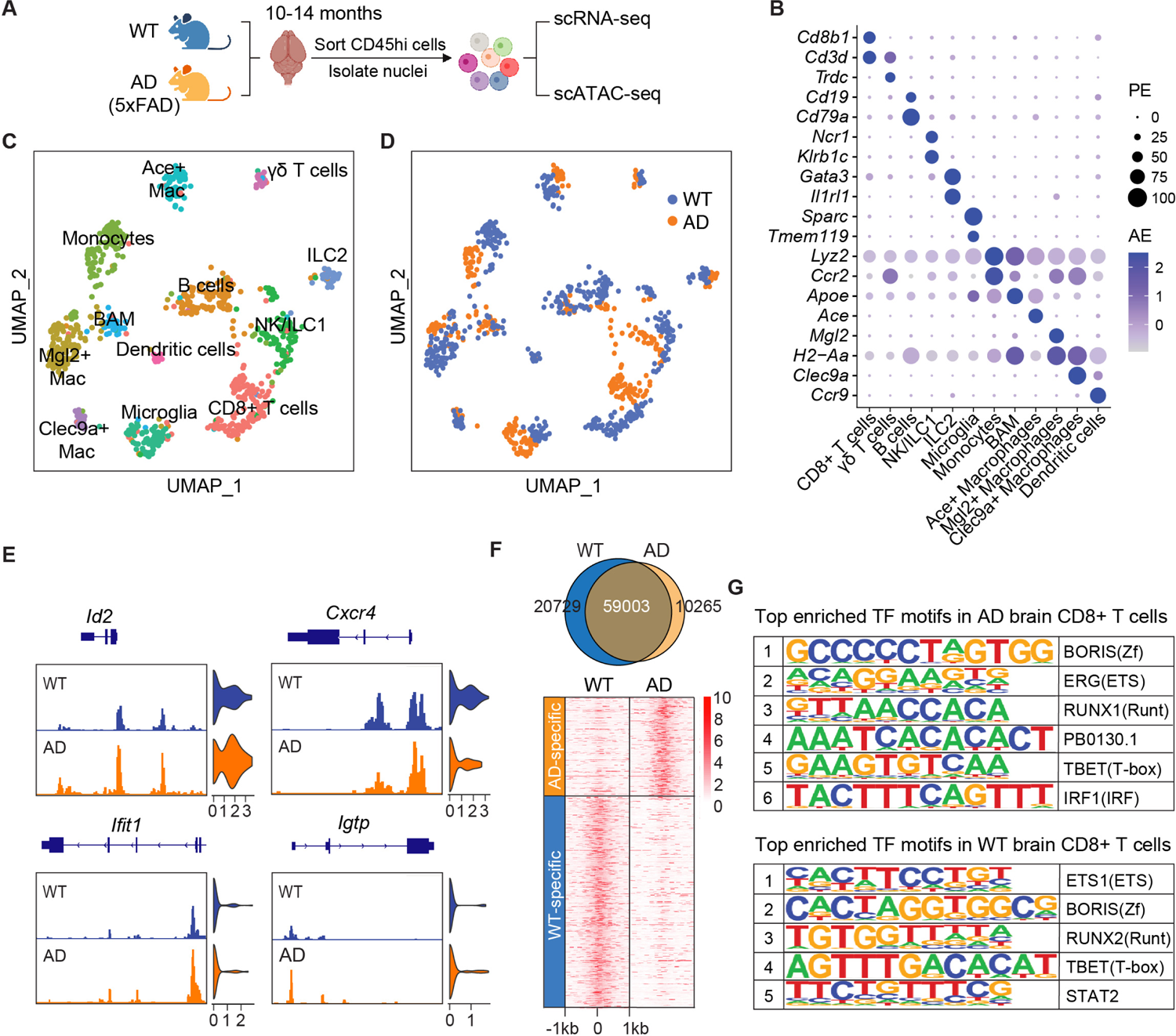
Multiomic profiling of lymphocytes in AD mouse brains identifies transcription factors (TF) that shape distinct regulome architecture. A. Experimental pipeline for multiomics analysis of wild type (WT) and 5xFAD (AD) brains at 10-14 months of age. CD45^hi^ cells were sorted, nuclei were isolated and were processed for scRNA-seq and scATAC-seq analysis on the same cells. B. Dot plot showing the expression of lineage genes for identification of UMAP clusters. C. UMAP showing the clustering of CD45^hi^ cells from WT (n=9, 516 cells) and AD (n=9, 341 cells) mouse brains. Mac, macrophages; BAM, border-associated macrophages. D. UMAP showing clustering of WT nuclei in blue, and AD nuclei in orange. E. IGV views of ATAC peaks at *Id2*, *Cxcr4*, *Ifit1* and *Igtp* loci in CD8+ T cells from AD mice compared with WT. Enriched peaks are found in *Cxcr4* in WT mice compared with AD. F. TF motifs enriched at differentially accessible regions (DARs) in CD8+ T cells. G. Top enriched TF motifs in CD8+ T cells from WT and AD mouse brains using HOMER.

We then aimed to identify transcription factors (TFs) in CD8+ T cells that altered chromatin profiles by analyzing TF motifs enriched at differentially accessible regions (DARs). Notably, there were two times more DARs specific to WT (n=20729) than AD (n=10265) CD8+ T cells (Figure 4F), consistent with more downregulated genes in the CD8+ T cells from AD mice (Supplementary Figure 3F). *De novo* TF motif enrichment analysis by HOMER showed the top enriched TF motifs in CD8+ T cells included common TF families of ETS, RUNX, T-box and Zf (Figure 4G). However, interferon regulatory factor 1 (IRF1), was amongst the top TF motifs specifically enriched in the AD-specific accessible regions, consistent with an upregulation of Type I IFN genes. Supporting this, we also found that a suite of other IRF motifs (ISRE, IRF8, IRF3, IRF2) were enriched in CD8+ T cells from AD mice using TF motif enrichment analysis by HOMER (Supplementary Figure 4). Interestingly, STAT2, a signal-dependent TF downstream of Type I IFN, was enriched in the WT-specific accessible regions, suggesting that Type I IFN signals also suppressed gene expression epigenetically. Together, these data indicate that the Type I IFN response in AD mice is both transcriptionally and epigenetically regulated.

## Enrichment of ISG+ CD8+ T cells in the brain

To assess systemic changes in the AD mice, we also conducted HDFC of CD45^hi^ cells from liver and spleen tissues of 10-12-month-old WT and AD mice and compared them with those from brain tissue (Figure 5). Using the Bioconductor CyTOF workflow, we performed multi-dimensional scaling (MDS) analysis of the HDFC data and found no substantial differences in liver and spleen CD45^hi^ cells between WT and AD mice (Figure 5A). By contrast, CD45^hi^ cells from the brain showed significant differences between WT and AD mice. We also compared the number of CD8+ T cells from the brain and outer dorsal meninges (dura mater and arachnoid mater) and found a localized increase within the AD mouse brain parenchyma, but not in the meninges (Figure 5B).

**Figure 5.**
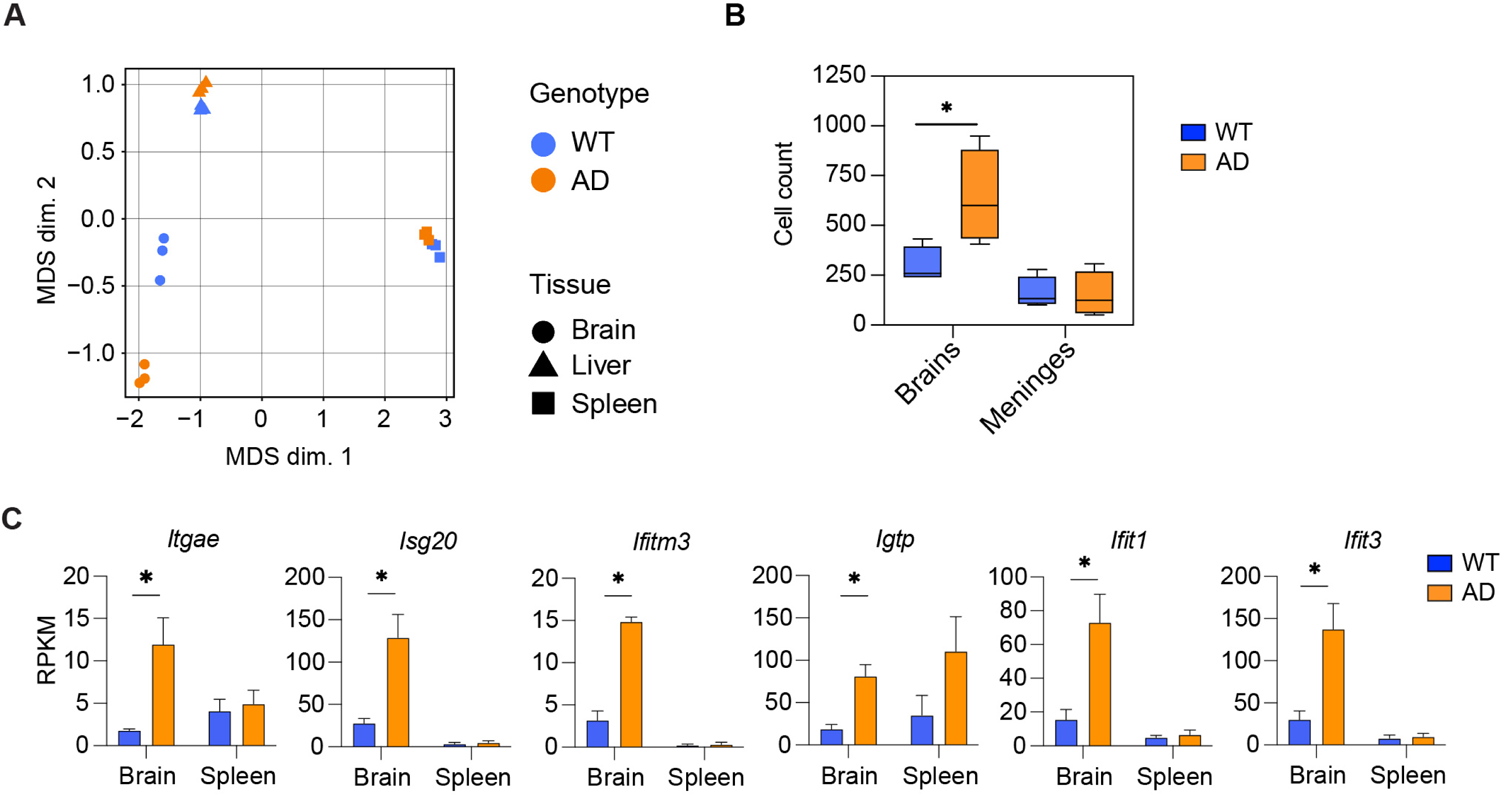
Enrichment of CD8+ T cells in AD in the brain. A. High-dimensional flow cytometry data from CD45^hi^ cells at 10-12 months in brain, liver and spleen. The multidimensional scaling (MDS) plot shows less distance between WT and AD groups in liver and spleen, but increased distance between WT and AD in the brain (n=3). B. Comparison of CD8+ T cell number between the brain and dorsal meninges shows a significant increase in AD mouse brains (p<0.05), but not in meninges (n=4). C. Low-input bulk RNA-seq on sorted CD8+ T cells from the brains and spleens of WT and AD mice at 10-14 months of age (n=3-4). Significantly higher gene expression (Reads Per Kilobase of transcript, per Million mapped reads, RPKM) of *Itgae*, *Isg20*, *Ifitm3*, *Igtp*, *Ifit1* and *Ifit3* genes were found in AD mouse brains (p<0.05), but not in spleen. Statistical analysis was performed using unpaired Student’s t tests. * denotes significance of p<0.05.

To further validate that the ISG+ CD8+ T cell transcriptomes were tissue-specific, we performed low-input bulk RNA-seq on sorted CD8+ T cells from the brains and spleens of WT and AD mice at 10-14 months of age. We found a significant upregulation of ISGs including *Itgae*, *Isg20*, *Ifitm3*, *Igtp*, *Ifit1*, and *Ifit3* only in the brains of AD mice in comparison to WT, but not in the spleen (Figure 5C). Taken together, the data suggest that the local, tissue-resident CD8+ T cell population within the brain parenchyma is transcriptionally distinct from peripheral CD8+ T cells.

## Blocking CD8+ T cell development attenuates memory loss and reduces amyloid plaques in AD mice

To investigate the impact of CD8+ T cells on memory impairment in AD mice, we utilized 5xFAD mice on a *B2m*-deficient background to inhibit MHCI expression and prevent the development of CD8+ T cells in the thymus of 5xFAD mice (Figure 6A). We then performed a Y-maze spontaneous alternation test to assess spatial working memory in the mice, which measures cognitive deficits by quantifying the mouse’s willingness to explore new environments. Normally, mice prefer to explore a new arm of the maze rather than revisiting one that has been previously explored. The AD mice demonstrated a lower frequency of spontaneous alternation than WT mice at both 4 and 6 months of age (Figure 6B-C). However, these cognitive deficits did not emerge at both 4 and 6 months in AD mice crossed with the *B2m*-deficient background that lacked CD8+ T cells. Given that CD8+ T cells exhibited ISG expression in our single cell multiomic data, we hypothesized these cells act through Type I IFN signaling. To confirm our hypothesis, we used 5xFAD mice on an *Ifnar1*-deficient background. Indeed, we observed the prevention of memory loss in these mice at both 4 and 6 months of age (Figure 6B-C), in line with the recent findings of the Cao lab using anti-IFNAR1 antibodies (Roy *et al*., 2022). We then stained brain slices for the expression of Aβ, in 5xFAD mice on a *B2m*-deficient background at 12 months of age (Figure 6D). Importantly, we found that 5xFAD-*B2m*^-/-^ showed significantly less Aβ plaques in the cortex, hippocampus, basolateral amygdala and thalamus regions, compared with 5xFAD mice (Figure 6D-E). Taken together, our data suggest that both CD8+ T cells and Type I IFN signaling are necessary to induce cognitive decline in AD, potentially operating through the same axis. Blocking CD8+ T cell development alleviates memory loss and significantly reduces Aβ plaques in AD mouse brains.

**Figure 6.**
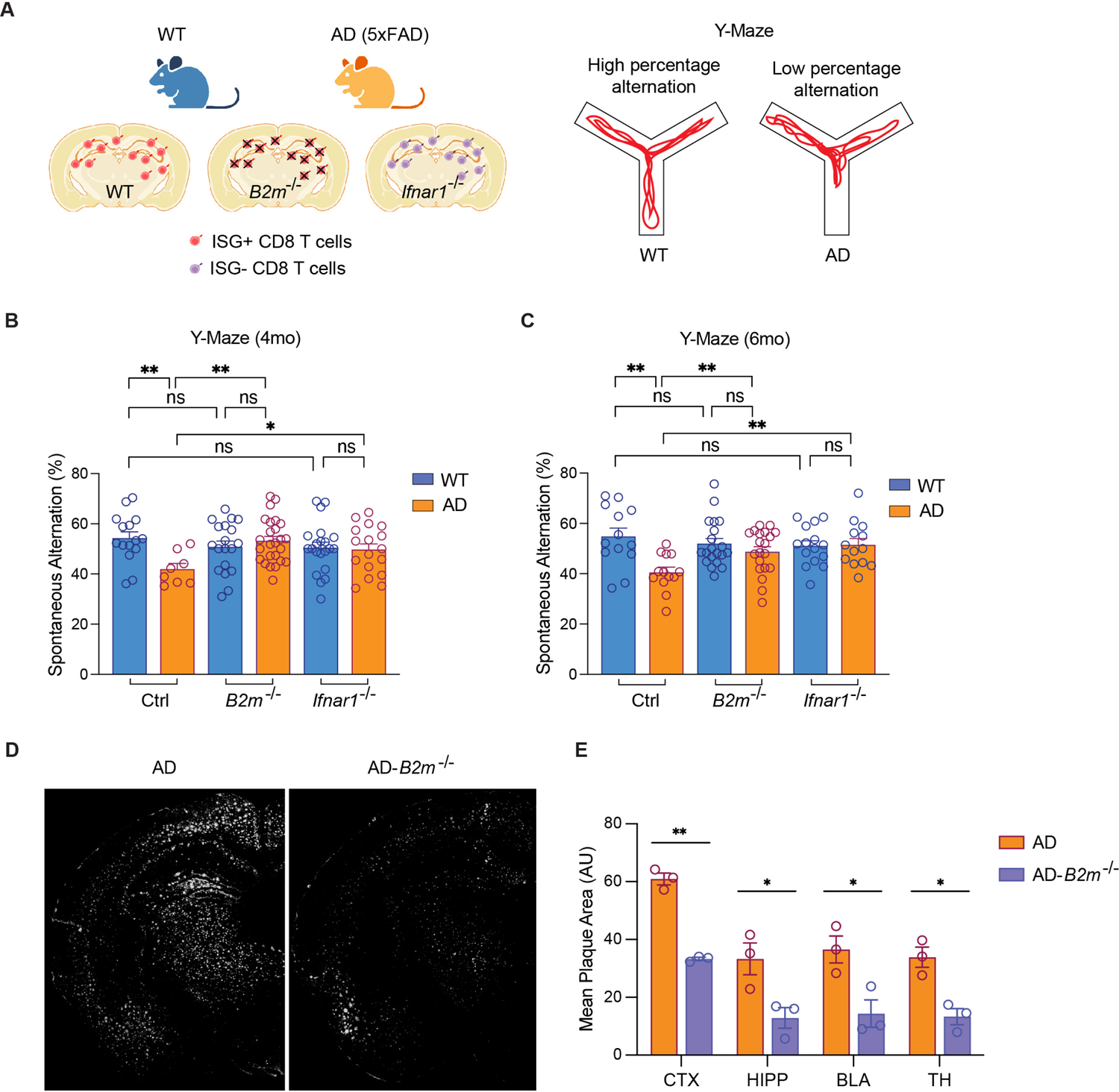
Blocking CD8+ T cell development rescues memory loss in AD mice. A. Schematic diagram depicting the three genetic backgrounds of mice used: C57BL/6J, *B2m*^-/-^ and *Ifnar1*^-/-^ mice. B-C. Y-maze spontaneous alternations in 4-and 6-month-old wild type (WT) mice (n=13-15), 5xFAD (AD) mice (n=8-13), WT-*B2m*^-/-^ mice (n=19-20), AD-*B2m*^-/-^ (n=21-25), WT-*Ifnar1*^-/-^ (n=15-21), and AD-*Ifnar1*^-/-^ (n=13-17). D. Representative immunohistochemical (IHC) images depicting amyloid-β (Aβ) staining in one hemisphere of coronal slices (40 µm thickness) from AD and AD-*B2m*^-/-^ mouse brains at 12 months of age. E. Mean Aβ plaque area (arbitrary units) in the cortex (CTX), hippocampus (HIPP), basolateral amygdala (BLA) and thalamus (TH) of AD and AD-*B2m*^-/-^ mouse brains (n=3). Statistical analysis was performed using unpaired Student’s t tests. * denotes significance of p<0.05. ** denotes significance of p<0.01. ns denotes no significance. AU, arbitrary units.

## Discussion

Although there is growing evidence of the immune system’s involvement (especially microglia) in neurodegenerative diseases, the precise signaling mechanisms of immune cells, including their regulatory networks and cytokine pathways, are still not understood (Chen and Colonna, 2022; Marsh *et al*., 2016; Mittal *et al*., 2019; Unger *et al*., 2020). To address this knowledge gap, we examined the transcriptomes and epigenomes of non-microglial immune cells in 5xFAD mice at various ages. Our results demonstrated that 1) tissue-resident CD8+ T cells were significantly enriched in aged AD mouse brains compared with young AD or aged WT brains; 2) a disease-associated sub-population of CD8+ T cells (DATs) that uniquely express high levels of ISGs was present at higher numbers in aged AD mouse brains; 3) CD8+ T cells had a distinct regulome in AD mice, and showed epigenomic control of Type I IFN signaling; 4) elimination of CD8+ T cells (*B2m*^-/-^) leads to a reduction in Aβ levels and improved spatial working memory in AD mice; and 5) elimination of Type I IFN signaling (*Ifnar1*^-/-^) also improves spatial working memory in AD mice. Together, our results indicate that DATs play a critical role in the pathogenesis of AD, potentially through Type I IFN signaling.

Our results highlight a tissue-resident transcriptional profile in aging AD mouse brains, including expression of CD103 (encoded by *Itgae*), which has been shown to be a key marker of tissue-resident CD8+ T cells that patrol the healthy human brain (Smolders *et al*., 2018). Interestingly, although CD8+ T cells have been proposed to interact with oligodendrocytes and microglia in aged and AD brains (Kaya et al., 2022; Marsh *et al*., 2016), our data suggest that in AD mice, a high frequency of CD8+ T cells were located in close proximity with neurons, potentially allowing for increased interactions. In addition, we have localized these resident CD8+ T cells within the mouse AD brain parenchyma, including the hippocampus and cortex, which supports previous findings in other AD models (Ferretti et al., 2016; Unger et al., 2020). These CD8+ T cells were largely absent in the dorsal meninges. Most recently, the Holtzman lab found that in models of tauopathy, both CD8+ and CD4+ T cells were increased in areas of tau pathology in 9.5-month-old mice, but interestingly not in age-matched mice with Aβ deposition (Chen et al., 2023). Further, they showed that T cell depletion ameliorated tau-mediated pathology and rescued memory loss (Chen *et al*., 2023).

Here, we critically demonstrate that memory loss was alleviated (at 4 and 6 months), and plaque deposition was reduced (at 12 months) in 5xFAD mice on a *B2m*-deficient background. A previous study showed that depletion of CD8+ T cells in APP/PS1 AD mice, using a systemically-delivered antibody, had no effect on cognitive function or plaque deposition (Unger *et al*., 2020), which is the opposite of our observations using *B2m*-deficient mice to block CD8+ T cell development. We hypothesize that treatment of AD mice with antibodies at a late stage of the disease may not be effective. The *B2m*-deficient model indicates that the elimination of CD8+ T cell signaling at early developmental stages may be required to influence spatial working memory. Depletion experiments using anti-CD8a delivered into the brain at an early stage of disease, or conversely, reconstitution of CD8+ T cells into *B2m*-deficient mice, may shed light on the potential of targeting tissue-resident CD8+ T cells as a treatment strategy. It is possible that these CD8+ T cells may also communicate with other cell types found to have an upregulation of MHCI genes, including disease-associated oligodendrocytes and microglia (Kaya *et al*., 2022; Kenigsbuch *et al*., 2022).

In the current study, we revealed an enrichment of a specific CD8+ T cell subset that express an antiviral-like Type I IFN-responding gene signature, here named as DATs. Type I IFN signaling is emerging as a key pathway involved in AD (Roy et al., 2020). Type I IFN signaling, and an expansion of CD8+ T cells also occur in other neurodegenerative diseases, including amyotrophic lateral sclerosis (ALS) (Campisi et al., 2022). These mechanisms should be studied further in a wider variety of neurodegenerative models, to establish the role of CD8+ T cell-specific Type I IFN signaling in neurodegenerative diseases, and identify which cells produce Type I IFNs (IFN-α and IFN-β) which may interact with CD8+ T cells. High-definition spatial transcriptomics methods may be useful in identifying these interactions. Our findings support a recent investigation in APP/PS1 mice showing that resident CD8+ T cells contribute to AD pathology through IFN-β signaling (Altendorfer *et al*., 2022). Another recent finding showed that there is an increase in ISG15+ CD8+ T cells in mouse brains with tauopathy. Further, our results implicate a role for Type I IFN signaling in memory loss in AD. We hypothesize that CD8+ T cell-specific Type I IFN signaling may be a critical contributor to this memory loss and should be further investigated.

Although Type I IFN signaling was enriched in the tissue-resident CD8+ T cells of AD mouse brains in our study, the complete role of these cells remains unclear. IFN-γ is a Type II IFN that is produced in high levels by CD8+ T cells, and can antagonize Type I IFN signaling (Platanias, 2005). This may explain why IFN-γ remained unchanged in CD8+ T cells from AD mice, even though *Ifng* was expressed in the ISG+ CD8+ T cell cluster. In other studies, the production of IFN-γ by infiltrating Aβ-specific Th1 cells in APP/PS1 mice promoted microglial activation, increased plaque burden, and was associated with cognitive impairment (Browne et al., 2013). Interestingly, inhibiting IFN-γ in mice with tauopathy reduced degeneration (Chen *et al*., 2023). Further studies using *Ifng*-deficient AD mice may shed light on the role of IFN-γ in AD, and whether global IFN signaling mechanisms are important in AD.

We also found that other cytokines and cytokine receptors, including chemokines, may participate in AD disease pathology. We identified that *Cxcr6* was upregulated in resident CD8+ T cells across all time points in AD mouse brains compared with WT. A recent investigation has shown that the CXCL16-CXCR6 axis is critical for monocyte interaction with clonally-expanded CD8+ T cells in the CSF of patients with cognitive impairment (Piehl *et al*., 2022). The increased availability of CXCL16 in the CSF, as well as the upregulation of *CXCR6* in CD8+ T cells, points to the CXCL16-CXCR6 axis as a key mechanism of entry for antigen-specific T cell entry into the brain. In our study, it is unclear if CD8+ T cells enter the brain parenchyma through the mouse CSF, and where potential expansion may occur. The choroid plexus epithelium forms the barrier between the brain parenchyma and the CSF and has been identified as a source of Type I IFN signaling, as well as a key site of lymphocyte infiltration (Suzzi et al., 2023). Investigation of cytokine signaling networks and their role in T cell entry into the AD brain are needed in future studies. We also found that the CCL5-CCR5 axis could be involved in aging, with decreased *Ccl5* and increased *Ccr5* found in aging brains. Interestingly, in patients with age-related macular degeneration, increased CCR5 expression in circulating CD8+ T cells was associated with slower atrophic lesion development (Krogh Nielsen et al., 2020).

In conclusion, we uncovered the transcriptional networks and functions of lymphocytes in an AD model and found that a tissue-resident Type I IFN-responding CD8+ T cell cluster was enriched in aged AD mouse brains. Importantly, we showed that CD8+ T cell (MHCI) deletion improved spatial working memory in AD mice and reduced amyloid plaque deposition. Together, our data point to a critical role for CD8+ T cells and Type I IFN signaling in the pathogenesis of AD. Further studies should examine cell-cell communication and cytokine-cytokine receptor networks between CD8+ T cells and other cell types in the brain, identify the lineage-specific mechanism of CD8+ T cell entry into the brain, as well as find the cellular source of Type I IFNs in AD. Strategies to block Type I IFN signaling and actions of tissue-resident CD8+ T cells may be useful in treating AD; however, our data suggest that early intervention may be required.

## Lead Contact

Further information and requests for resources and reagents should be directed to and will be fulfilled by the Lead Contact, Han-Yu Shih.

## Data and Code Availability

The low input RNA-seq, scRNA-seq and single cell multiome data have been deposited in the GEO: GSE227130 (token upon request).

## Experimental Model and Subject Details

### Animals

All animals were maintained in the AAALAC-accredited animal housing facilities at the National Institutes of Health (NIH) on a 12/12hr light/dark cycle. All animal studies were performed according to NIH guidelines for the use and care of live animals and were approved by the Institutional Animal Care and Use Committee of the National Eye Institute (NEI). All mice were either genotyped manually according to pre-determined primer/probe sets (The Jackson Laboratory) or were submitted for automated genotyping PCR services (Transnetyx, Cordova, TN).

Male and female 5xFAD heterozygous transgenic mice on a B6 background (#034848, B6.Cg-Tg (APPSwFlLon,PSEN1*M146L*L286V)6799Vas/Mmjax, The Jackson Laboratory, Bar Harbor, ME) were used throughout this study. 5xFAD is a transgenic model that co-expresses 5 mutations in APP and PSEN1 to reproduce pathological features of AD in mice (Oakley *et al*., 2006). Sex-and age-matched littermate non-carrier wild type mice (WT, or +/+), or B6 mice were used as controls during the experiments.

We also utilized hAbeta^SAA^ knock-in (APP-SAA KI) mice, which introduced 3 humanizing Aβ region mutations to endogenous exons without disturbing intrinsic gene expression (Xia et al., 2022). Male and female APP-SAA KI homozygous transgenic mice on a B6 background (#034711, The Jackson Laboratory) were also used in this study. Sex-and age-matched littermate non-carrier wild type mice (WT, or +/+) were used as controls during the experiments.

5xFAD mice crossed with *B2m*-deficient mice (#002087, The Jackson Laboratory) were used to block CD8+ T cell development. 5xFAD mice crossed with *Ifnar1*-deficient mice (# 028288, The Jackson Laboratory) were used to create AD mice without functional Type I IFN signaling.

## Tissue dissection and processing

Mice were deeply anesthetized using 10% v/v Euthanasia solution (VetOne, Boise, ID) with 2% v/v Xylocaine (Fresenius Kabi, Bad Homburg, Germany) in 1x phosphate buffered saline (PBS). Mice were then perfused with 20 ml 1x PBS through transcardial perfusion. Spleen, liver, brain and/or meninges were then dissected from each mouse for processing into a single cell suspension, and collected into RPMI 1640 (#11875119, Thermo Fisher Scientific, Waltham, MA). For regional brain studies, whole brains were dissected to isolate the cortex, hippocampus or thalamus according to previously published methodology (Farías et al., 2016).

Spleens were mashed through a 70 µm filter (#1181X55, Thomas Scientific, Swedesboro, NJ) and were centrifuged at 1500 rpm for 5 minutes at 4°C. Samples were resuspended in 1 ml RPMI and were counted using a Cellometer Auto 2000 and ViaStain AOPI Staining Solution (Nexcelom Bioscience, Lawrence, MA). 1 million splenocytes were used for further staining for flow cytometry or sorting.

Livers and brains were minced in RPMI using a scalpel blade (#72042-11, Electron Microscopy Services, Hatfield, PA) and then were incubated with 2.5 mg/ml Collagenase IV (#LS004189, Worthington Biochemicals, Lakewood, NJ) and 0.1 mg/ml DNAse I (#10104159001, Sigma-Aldrich, St. Louis, MO) for 30 minutes at 37°C with shaking at 200 rpm. Livers and brains were then mashed and rinsed through a 70 µm filter with RPMI on ice and were centrifuged at 1500 rpm for 5 minutes at 4°C. Samples were resuspended in 6 ml RPMI and were mixed with 4 ml 90% Percoll (diluted in 10x PBS, #GE17-0891-01, Sigma-Aldrich) at room temperature. Only brain samples were then layered carefully over 2 ml 70% Percoll at room temperature to produce a density gradient. All samples were centrifuged at 1500 rpm for 30 minutes at 18°C with minimal braking speed to not disturb the gradient. Brain cells at the gradient interphase were collected and washed in 1x PBS. All brain cells collected were used for further staining. Liver cells were resuspended in 1 ml RPMI and were counted using a Cellometer Auto 2000 and ViaStain AOPI Staining Solution. 1 million liver cells were used for further staining.

Meninges were dissected from the skullcaps under a stereo dissecting microscope (#SZ61, Olympus, Tokyo, Japan) using previously published methodology (DiSano et al., 2020; Louveau et al., 2018). Meninges were collected into RPMI and were incubated with 2.5 mg/ml Collagenase IV and 0.1 mg/ml DNAse I for 30 minutes at 37°C with shaking at 300 rpm. Meninges were pipetted up and down until a single cell suspension was observed. Cells were then filtered through a 70 µm filter top tube (#352235, Thermo Fisher Scientific) and were centrifuged at 1500 rpm for 5 minutes at 4°C. All meningeal cells collected were used for further staining.

## Preparation of single cell suspensions for flow cytometry and cell sorting

Cells were transferred to a 96-well round bottom plate (#29442-066, VWR, Radnor, PA) and were washed twice with wash buffer (1x PBS containing 1% fetal bovine serum, FBS). Cells were incubated with a staining and blocking cocktail for 20-30 minutes at 4°C under dim conditions. Following antibody staining, cells were washed twice with wash buffer and were resuspended in wash buffer for flow cytometry or cell sorting.

Cells were analyzed on a Cytek Aurora (5-laser, 40-color instrument) with SpectroFlo software (Cytek, Fremont, CA) at the National Institute of Arthritis and Musculoskeletal and Skin Diseases (NIAMS Flow Cytometry Section). 60K events were acquired per brain sample for comparison. Flow cytometry data were processed and analyzed using FlowJo v9 (FlowJo, LLC; BD, Franklin Lakes, NJ), Bioconductor CyTOF workflow (Nowicka et al., 2022) and GraphPad Prism 8 (GraphPad, San Diego, CA) software. For cell sorting, cells were analyzed and sorted on a FACS Aria Fusion (BD) and were collected into RPMI containing 10% FBS for downstream analysis.

For lymphocyte profiling on the Cytek Aurora, the following staining panel was used: Fc blocking (anti-CD16/32, clone 93, 1:100), LIVE/DEAD Fixable Blue Stain (1:1000), anti-CD11b (clone M1/70, 1:200), anti-CD19 (clone 6D5, 1:500), anti-CD4 (clone RM4-5, 1:400), anti-CD44 (clone IM7, 1:200), anti-CD45 (clone 30-F11, 1:200), anti-CD49a (clone Ha31/8, 1:400), anti-CD49b (clone HMa2, 1:800), anti-CD62L (clone MEL-14, 1:800), anti-CD69 (clone H1.2F3, 1:100), anti-CD8a (clone 53.6.7, 1:200), anti-KLRG1 (clone 2F1/KLRG1, 1:400), anti-NK1.1 (clone PK136, 1:100), anti-NKp46 (clone 29A1.4, 1:100), anti-ST2 (clone U29-93, 1:100), anti-TCRb (clone H57-597, 1:200) and anti-TCRd (clone GL3, 1:50).

For sorting on the BD FACS Aria Fusion for CD45^hi^ cells, Fc blocking (anti-CD16/32, clone 93, 1:100), LIVE/DEAD Fixable Blue Stain (1:1000) and anti-CD45 (clone I3/2.3, 1:200) were used. For CD8+ T cell sorting, the following staining panel was used: Fc blocking (anti-CD16/32, clone 93, 1:100), polyclonal F(ab’)₂ fragment for blocking (1:100), FBS (1:20), LIVE/DEAD Fixable Blue Stain (1:1000), anti-CD11b (clone M1/70, 1:200), anti-CD19 (clone 6D5, 1:500), anti-CD11c (clone HL3, 1:200), anti-TER119 (clone TER-119, 1:200), anti-CD45 (clone 30-F11, 1:200), anti-TCRb (clone H57-597, 1:100), anti-CD4 (clone RM4-5, 1:200) and anti-CD8a (53-6.7, 1:200).

## Single cell RNA-sequencing

Cell hashing reagents (TotalSeq anti-mouse hashtags B0301-306, BioLegend) were used to tag cells from biological replicates prior to pooling and sorting, according to the manufacturer’s protocol. Brain CD45^hi^ cells were sorted from each genotype using a FACS Aria Fusion (BD). ∼20K cells at a concentration of 800 cells/µl were processed using the Chromium Next GEM Single Cell 3ʹ GEM, Library & Gel Bead Kit v3.1 (10x Genomics, #1000121), Chromium Next GEM Chip G Single Cell Kit (10x Genomics, #1000120) and Chromium Controller (10x Genomics, #1000202). cDNA and cell surface protein libraries were constructed using the Chromium Single Cell 3ʹ Feature Barcode Library Kit (10x Genomics, #1000079) and Single Index Kit T Set A (10x Genomics, #1000213). cDNA libraries were sequenced at ∼500 million reads per sample with 50bp paired-end reads using NovaSeq or NextSeq (Illumina, San Diego, CA). All sequencing was performed in the NIAMS Genomic Technology Section.

Sequencing reads were demultiplexed and mapped to the mm10 reference transcriptome using Cell Ranger Single Cell v7.0 software (10x Genomics) to generate the FASTQ files and gene expression and feature barcode counts. The matrix files generated from Cell Ranger were further analyzed using Seurat v4.3.0(Satija et al., 2015) for quality control and filtering, normalization, clustering, visualization and differential gene expression analysis of cell populations. Gene Ontology (GO) analysis were performed using Enrichr (Chen et al., 2013) and ggplot2 (Wickham, 2016) in R v4.2.2. Downstream analyses were performed with custom R programs (R Core Team, 2022).

## Single cell multiomics analysis

Brain CD45^hi^ cells were sorted from each genotype using a FACS Aria Fusion (BD). Cells were permeabilized and nuclei were washed and quantified according to 10x Genomics protocols. ∼20K cells per sample were used for preparation of the nuclei for the Assay for Transposase-Accessible Chromatin (ATAC) and gene expression libraries using the low cell input nuclei isolation protocol (10x Genomics). The Chromium Next GEM Single Cell Multiome ATAC + Gene Expression Reagent Bundle (10x Genomics, #1000283), Chromium Next GEM Chip J Single Cell (10x Genomics, #1000234), Dual Index Kit TT Set A (10x Genomics, #1000215), Single Index Kit N Set A (10x Genomics, #1000212) and Chromium Controller were used according to the manufacturer’s protocols. For the ATAC library, a sequencing depth of 25K read pairs per nucleus was used with 50bp paired-end reads using NovaSeq or NextSeq (Illumina). For the gene expression library, a sequencing depth of 20K read pairs per nucleus was used. All sequencing was performed in the NIAMS Genomic Technology Section.

Sequencing reads were processed using Cell Ranger 7.0 (10x Genomics) to generate the FASTQ files, gene expression and ATAC combined counts. The matrix files generated from Cell Ranger were further analyzed using Seurat v4.3.0 and Signac (Satija *et al*., 2015; Stuart et al., 2021) to simultaneously measure the transcriptome and chromatin accessibility from the same cell. RNA counts were processed independently by quality control filtering, normalization, clustering, and differential gene expression. ATAC counts were processed independently by quality control filtering, macs2 peak calling, normalization, and clustering. Finally, an integrative joint analysis of RNA and ATAC data from each cell was performed using ‘Weighted Nearest Neighbor’ (WNN) methods in Seurat v4.3.0 to compute a joint neighbor graph and joint clustering that represents both gene expression and chromatin accessibility measurements from the same cell. The joint clustering was followed by differential peak analysis, peak annotation and linking peaks to genes by computing the correlation between the gene expression and chromatin accessibility at nearby peaks. The peak to gene links were then visualized using coverage plots. Further downstream analyses were performed with custom R programs (R Core Team, 2022). IGV 2.3 was used to visualize ATAC peaks. Transcription factor motif analysis was conducted using HOMER.

## Low-input RNA-sequencing

Brain (2000-5000 cells per sample) and spleen (2000 cells per sample) CD8+ T cells were sorted for low-input RNA-seq. cDNA was synthesized, amplified and libraries constructed using the NEBNext Single Cell/Low Input RNA Library Prep Kit for Illumina (E6420, NEB, Ipswich, MA) and NEBNext Multiplex Oligos for Illumina (E6440, NEB), according to the manufacturer’s protocols. cDNA libraries were sequenced at 40 million reads per sample with 50bp paired-end reads using NovaSeq or NextSeq (Illumina). All sequencing was performed in the NIAMS Genomic Technology Section.

The run was demultiplexed and converted to FASTQ format using bcl2fastq v2.20.0.422 (Illumina) and mapped to mm10 using Tophat 2.1.0 (Trapnell et al., 2012). Partek Genomics Suite 7.0 (Partek Inc., St. Louis, MO) and custom R programs were used for downstream analyses.

## Histology and immunohistochemistry

Mice were perfused with 20 ml 1x PBS followed by 10 ml 4% paraformaldehyde. After euthanasia, brains were dissected and post-fixed in 4% paraformaldehyde for 24 hours. Brains were then transferred to 30% sucrose for another 24 hours prior to embedding in Tissue-Tek Optimal Cutting Temperature Compound (Sakura Finetek USA, Torrance, CA). Samples were embedded and cryosectioned at 10 µm in the coronal plane by Histoserv Inc. (Germantown, MD).

Multiplexing immunohistochemistry (IHC) was performed by the NINDS Flow and Imaging Cytometry Core Facility. The following primary antibodies were used (1:200): anti-CD4 (BioLegend, #100538), anti-IBA1 (Wako Chemicals, Osaka, Japan, #019-19741), anti-GFAP (Agilent/Dako, #Z0334), anti-NeuN (Millipore Sigma, #ABN90P), anti-Aβ (Novus Biologicals, Englewood, CO, #NBP2-13075) and anti-CD8a (Thermo Fisher Scientific, #14-0195-82). Secondary antibodies from Thermo Fisher Scientific were then used (1:500): Alexa Fluor 546 goat anti-mouse IgG1 (#A21123), Alexa Fluor 647 goat anti-mouse IgG2a (#A21241), Alexa Fluor 488 goat anti-mouse IgG2b (#A21141), Alexa Fluor 488 goat anti-rabbit (#A11008), Alexa Fluor 555 goat anti-guinea pig (#A21435), or Alexa Fluor 647 goat anti-rat (#A21247). Analysis of CD4+ and CD8+ T cells were performed by adding masks to the cell for counting (masks remain in the figure to aid in visualization).

For all other IHC staining, brains were sectioned in the coronal plane at 40 μm, blocked for 1 hour with 10% goat serum, probed for 1 hour against Aβ (1:2000, Novus Biologicals, #NBP2-13075), washed with PBS, incubated for 1 hour with Alexa Fluor 594 goat anti-mouse (1:500, Thermo Fisher Scientific, #A11032), and Hoechst stain for 5 minutes (1:6000, Thermo Fisher Scientific, #62249). Sections were mounted and imaged using the Mica widefield confocal microscope (Leica, Wetzlar, Germany) at 1000x (tiled and stitched). Analysis was performed using FIJI (ImageJ, NIH, Bethesda, MD) for the Mean Grey Area from four regions (retrosplenial cortex, hippocampus, basolateral amygdala, and thalamus) in 5xFAD and 5xFAD-*B2m*^-/-^ 12-month-old mice.

## Y-maze behavioral assay

The spontaneous alternation Y-maze was performed on WT, 5xFAD, WT-*B2m*^-/-^, 5xFAD-*B2m*^-/-^, WT-*Ifnar1*^-/-^ and 5xFAD-*Ifnar1*^-/-^ mice aged 4 and 6 months. The assay was conducted in a symmetrical tan Plexiglas Y-Maze with three arms (38 cm long × 7.5 cm wide × 12.5 cm high) at 120° angles, designated A, B, and C. The mice were placed in the distal end of arm A and allowed to explore the maze for 5 minutes. A video camera mounted above the maze recorded the movements of the mice for analysis. Using TopScan Suite 3.0 (CleverSys Inc., Reston, VA) the arm entries were recorded, and the percentage of alternations (entry into an arm that differs from the previous two entries) was calculated.

## Statistical analysis

All graphing was performed using FlowJo, GraphPad Prism 8 or RStudio. Statistical analysis was performed using GraphPad Prism 8. IHC images may be presented in pseudo colors for visualization purposes. Image panels were constructed using BioRender.com (Toronto, Canada) and Adobe Illustrator (San Jose, CA). Further information related to the statistical analyses performed and sample sizes is provided in the figure legends.

## Acknowledgements

We sincerely thank the following core facilities at NIH for their support of this work: NIAMS Flow Cytometry Section, Genomic Technology Section, and Light Imaging Section; NEI Flow Cytometry Core Facility, and the Animal Facility; NINDS Flow and Imaging Cytometry Core Facility; NIMH IRP Rodent Behavioral Core; and the NHLBI Flow Cytometry Core. This study utilized the high-performance computational capabilities of the Biowulf Linux cluster at the NIH. This work was supported by the Intramural Research Programs of NEI and NINDS at the NIH (ZIAEY000569-01).

## Author Contributions

N.F. and H-Y.S. designed the project and wrote the manuscript with advice and revisions from all authors. N.F., J.G., A.B., L.R., C.L., V.B., M.B., W.M., R.D-P., M.M., and D.M. performed and analyzed the experiments. V.N. and S.B. helped with genomic data processing and analyses. R.C. and D.B.M. provided intelligent input. H-Y.S. supervised the project.

## Supplementary Figure Legends

**Supplementary Figure 1, related to Figure 1.**
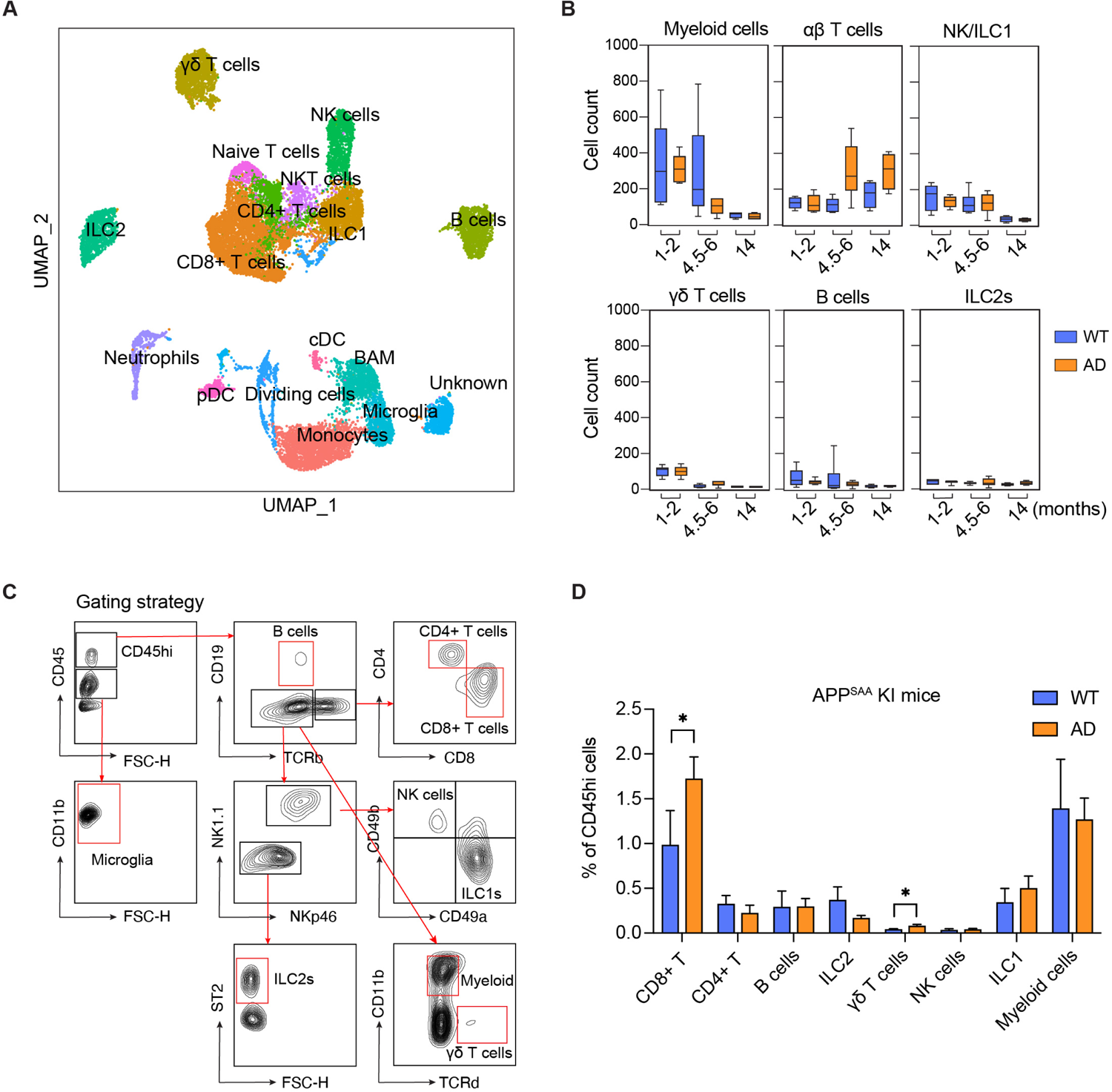
A. scRNA-seq UMAP of 20,874 cells combined between all experimental groups (n=4-6 pooled mice per group) showing detailed labeling of lymphocyte and myeloid cell clusters. 20,874 cells were pooled from wild type (WT) 1-2 months (5,014 cells), 5xFAD (AD) 1-2 months (4,505 cells), WT 4.5-6 months (3,829 cells), AD 4.5-6 months (3,731 cells), WT 10-14 months (1,677 cells) and AD 10-14 months (2,118 cells). B. Cell hashing data showing individual replicates of cell counts for myeloid cells, αβ T cells, NK/ILC1s, γδ T cells, B cells and ILC2s in WT and AD mice (n=4-6). B. Gating strategy used to separate lymphocyte populations using FlowJo. Live, singlet CD45+ cells were separated into CD45^hi^ (lymphocyte and myeloid cells) and CD45^int^ CD11b^hi^ (microglia). CD45^hi^ cells were further separated into B cells (CD19+, TCRb-), CD4+ T cells (CD19-, TCRb+ CD4+), CD8+ T cells (CD19-, TCRb+, CD8+), γδ T cells (CD19-, TCRb-, CD11b-, TCRd+), NK cells (CD19-, TCRb-, NK1.1+/NKp46+, CD49a-, CD49b+), ILC1s (CD19-, TCRb-, NK1.1+/NKp46+, CD49a+, CD49b-), ILC2s (CD19-, TCRb-, NK1.1-, NKp46-, ST2+) and myeloid cells (CD19-, TCRb-, CD11b+). C. Flow cytometry analysis (n=3) of CD45^hi^ cells from brains of WT and APP^SAA^ KI (AD) mice at 8-9 months of age. The graph shows the % of CD45^hi^ cells of B cells, CD4+ T cells, CD8+ T cells, γδ T cells, NK cells, ILC1s, ILC2s and myeloid cells. A significant increase in CD8+ T cells and γδ T cells in AD mice was found compared with WT at 8-9 months of age (p<0.05). Statistical analysis was performed using multiple unpaired Student’s t tests. * denotes significance of p<0.05.

**Supplementary Figure 2, related to Figure 2.**
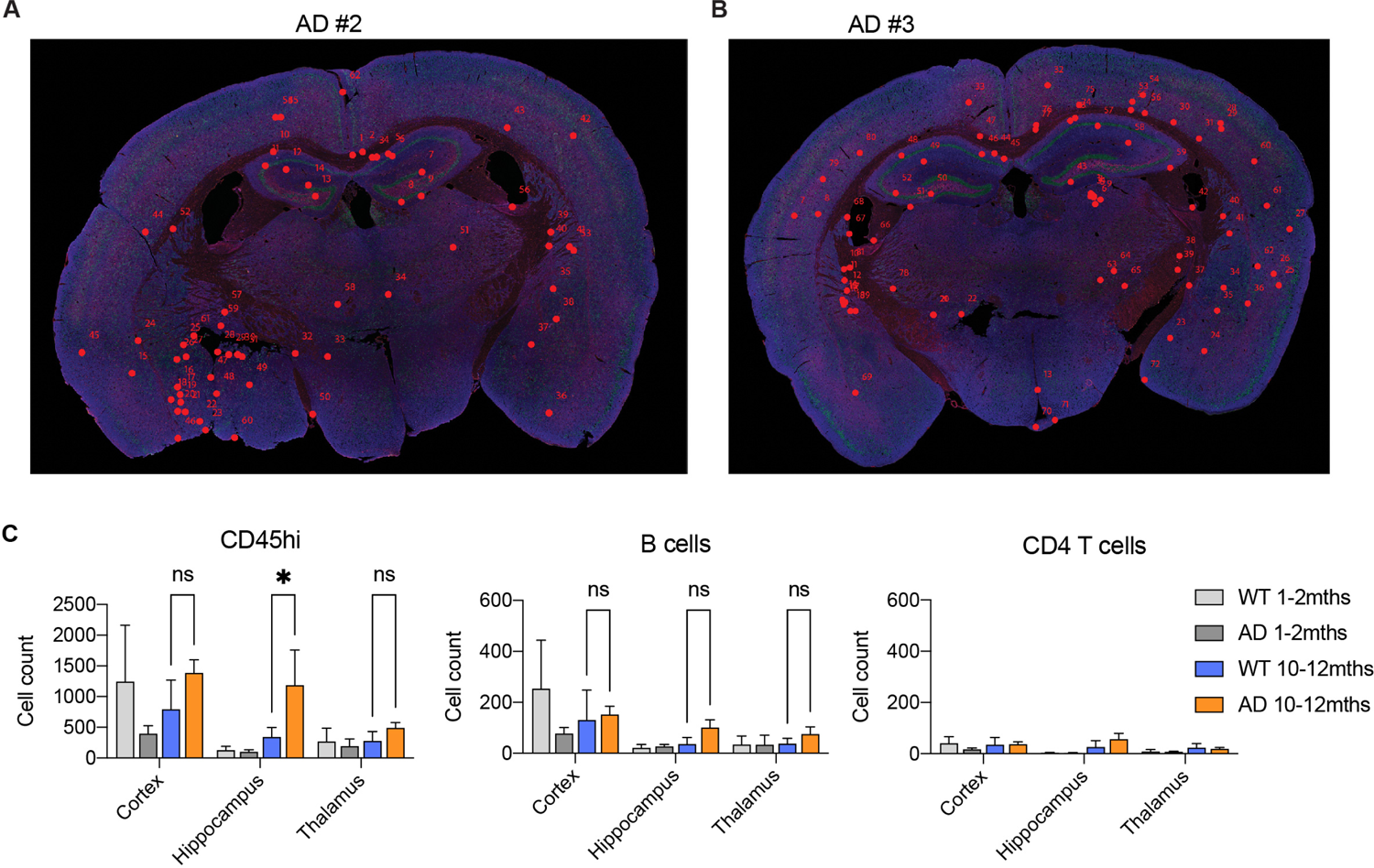
A-B. Localization of CD8+ T cells (red) in coronal brain slices of AD mouse #2 and AD mouse #3. C. Flow cytometry analysis of CD45^hi^ cells, B cells and CD4+ T cells in the cerebral cortex, hippocampus and thalamus of wild type (WT) and 5xFAD (AD) brains at 1-2 months and 10-11 months of age. Statistical analysis was performed using unpaired Student’s t tests. * denotes significance of p<0.05.

**Supplementary Figure 3, related to Figure 3.**
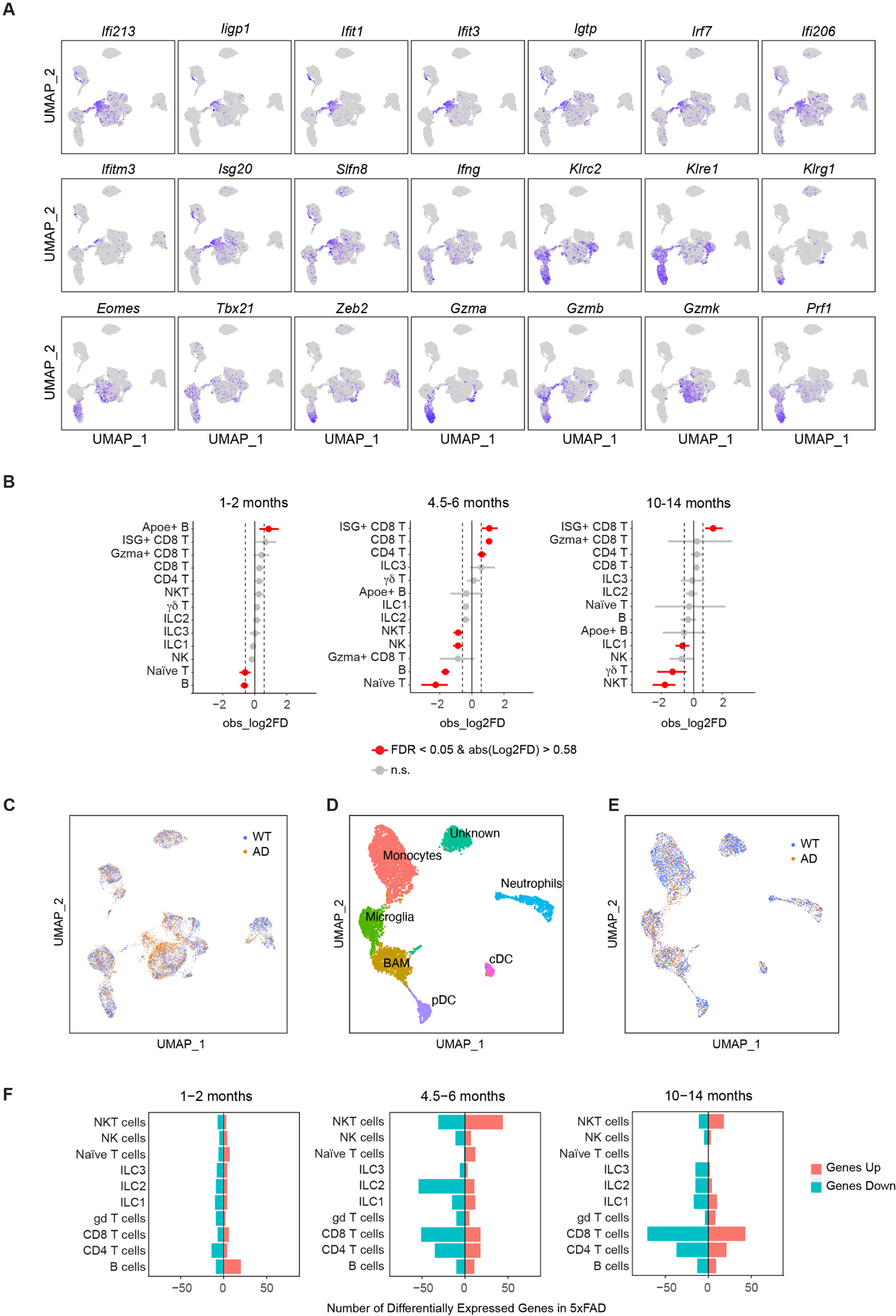
A. Feature plots showing the expression of IFN-stimulated genes (ISGs), *Irf7*, *Ifng*, effectors and transcription factors in lymphocytes. B. Single cell proportion tests of the 13 lymphocyte clusters at the 3 time points. Significant values are in red (q<0.05) and fold differences (FD) are set at −0.58 and 0.58. Statistical analysis was performed using the scProportionTest in R. C. UMAP of wild type (WT, blue) vs 5xFAD (AD, orange) showing enrichment in CD8+ T cell clusters in the AD mice. D. scRNA-seq UMAP of myeloid cells only depicting 7 cell clusters from WT and AD mouse brains at 1-2, 4.5-6 and 10-14 months of age. E. UMAP of WT (blue) vs AD (orange) showing no major difference in myeloid cell clusters in the AD mice. F. Bidirectional graphs showing the number of differentially expressed genes (DEGs) in AD mouse brains compared with WT at each time point (details of DEGs are provided in Supplementary Table 1). Upregulated genes are in red, downregulated genes are in blue. * denotes significance of p<0.05.

**Supplementary Figure 4, related to Figure 4.**
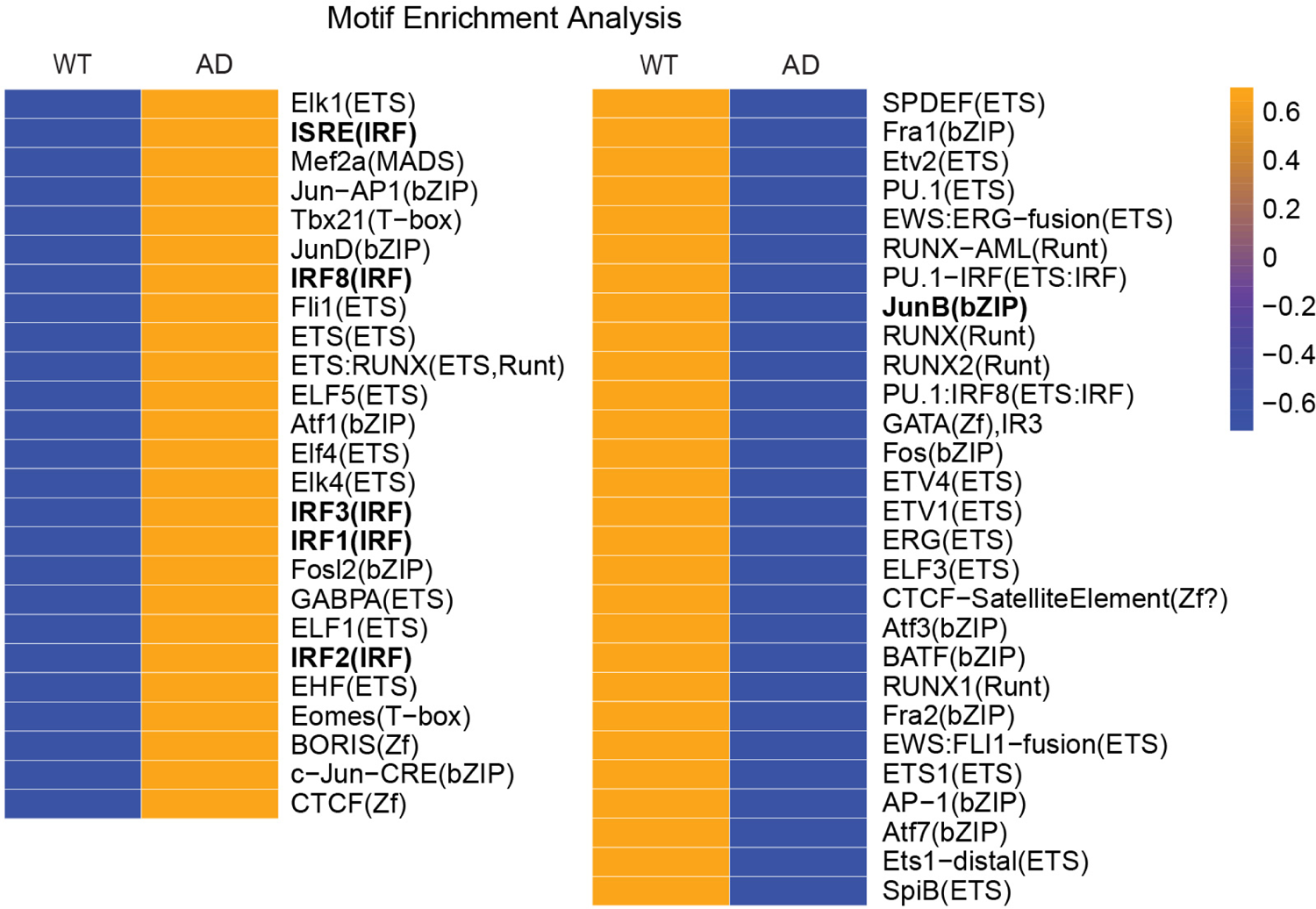
Known transcription factor (TF) motif enrichment analysis in wild type (WT)-and 5xFAD (AD)-specific accessible regions using HOMER. A higher frequency of motifs in CD8+ T cells is displayed in orange, and a lower frequency in blue.

## Notes

### Competing Interest Statement

The authors have declared no competing interest.

